# Genomic analysis unveils the role of genome degradation events and gene flux in the emergence and persistence of *S*. Paratyphi A lineages

**DOI:** 10.1101/2022.06.09.495420

**Authors:** Jobin John Jacob, Agila K Pragasam, Karthick Vasudevan, Aravind V, Monisha Priya T, Tharani Priya T, Pallab Ray, Madhu Gupta, Arti Kapil, Sulochana Putil Bai, Savitha Nagaraj, Karnika Saigal, Temsunaro Rongsen Chandola, Maria Thomas, Ashish Bavdekar, Sheena Evelyn Ebenezer, Jayanthi Shastri, Anuradha De, Shantha Dutta, Anna P Alexander, Roshine Mary Koshy, Dasaratha R Jinka, Ashita Singh, Sunil Kumar Srivastava, Shalini Anandan, Gordon Dougan, Jacob John, Gagandeep Kang, Balaji Veeraraghavan, Ankur Mutreja

## Abstract

Paratyphoid fever caused by *S*. Paratyphi A is endemic in parts of Asia and Sub-Saharan Africa. The proportion of enteric fever cases caused by *S*. Paratyphi A has substantially increased, yet only limited data is available on the population structure and genetic diversity of this serovar. We examined the phylogenetic distribution and evolutionary trajectory of *S*. Paratyphi A isolates collected as part of the Indian enteric fever surveillance study “Surveillance of Enteric Fever in India (SEFI).” In the study period (2017-2020), *S*. Paratyphi A comprised 17.6% (441/2503) of total enteric fever cases in India, with the isolates highly susceptible to all the major antibiotics used for treatment except fluoroquinolones. Phylogenetic analysis clustered the global *S*. Paratyphi A collection into seven lineages (A-G), and the present study isolates were distributed in lineages A, C and F. Our analysis documented that the genome degradation events and gene acquisitions or losses play a major role in the evolution of new *S*. Paratyphi A lineages/sub-lineages. A total of 10 pseudogene-forming mutations possibly associated with the emergence of lineages were identified. Pan-genome analysis identified the insertion of P2/PSP3 phage and acquisition of IncX1 plasmid during the selection in 2.3.2/2.3.3 and 1.2.2 genotypes, respectively. We also identified that the six characteristic missense mutations associated with the lipopolysaccharide (LPS) biosynthesis genes of *S*. Paratyphi A confer only a low structural impact and would therefore have minimal impact on vaccine effectiveness. Since *S*. Paratyphi A is human restricted, high levels of genetic drift are not expected unless these bacteria transmit to naive hosts. However, public-health investigation and intervention by means of genomic surveillance would be continually needed to avoid *S*. Paratyphi A serovar becoming a public health threat similar to the *S*. Typhi of today.

## Introduction

Enteric fever is a life-threatening systemic febrile illness caused by infections with *Salmonella enterica* serovar Typhi, Paratyphi A, B and C [1]. *S*. Typhi is the predominant cause of enteric fever, with an estimated 12 - 25 million cases of typhoid per year globally [2]. Among the three serovars that cause paratyphoid fever, *S*. Paratyphi A is the most prevalent and infections with *S*. Paratyphi B and C serotypes are extremely rare [3]. Both Typhoid and paratyphoid infections are endemic in parts of South-central Asia, South East Asia and Sub-Saharan Africa [4]. Though only limited data is available on the true burden of *S*. Paratyphi A in these regions, it is estimated to cause around 5 million cases of enteric fever annually [5]. However, the actual number of infections was underestimated as paratyphoid is clinically indistinguishable from typhoid fever [6]. Recent data suggests that the proportion of enteric fever cases caused by *S*. Paratyphi A has substantially increased from 20% to 50% in some endemic regions of South Asia [7].

The sequential emergence of antimicrobial resistance in serovar Typhi over the past 50 years is well documented. Clinical, laboratory and genomic features of the evolution of antimicrobial resistance in *S*. Typhi against chloramphenicol (1960), first-line antimicrobials (1990), fluoroquinolones, third-generation cephalosporins and azithromycin are already established [8 - 9]. However, unlike *S*. Typhi, serovar Paratyphi A is predominantly susceptible to most antibiotics. Nevertheless, high fluoroquinolone non-susceptibility in *S*. Paratyphi A has been witnessed in recent years, with sporadic reports of multidrug resistant (MDR) and azithromycin resistant isolates [10 - 11].

*S*. Paratyphi A was found to have substantial regional differences with the emergence of seven distinct lineages (A-G), each having originated in a specific geographical location [12]. Among the lineages, A and C have expanded throughout South Asia and Southeast Asian countries to become successful clones, whereas other lineages are still rare. Unlike *S*. Typhi, the genome-level difference of *S*. Paratyphi A was investigated in only a few isolates [13 - 14]. Interestingly, evolutionary changes in *S*. Paratyphi A by means of gene gain or loss or mutations are mostly considered transient and are continuously removed by purifying selection [12]. However, a positive selection that may favor the diversification and expansion of certain lineages has not been studied previously. Here, we examined the phylogenetic distribution of *S*. Paratyphi A isolates collected as part of the Indian enteric fever surveillance named Surveillance of Enteric Fever in India (SEFI). We also examined the gain, loss and inactivation of genes at the genomic level to shed light on the ongoing process of evolution in *S*. Paratyphi A.

## Results

### Surveillance of S. Paratyphi A infections

During the study period between October 2017 to September 2020, 441 *S*. Paratyphi A were isolated from blood and bone marrow cultures performed at all study sites. Laboratory-based surveillance in tertiary care hospitals yielded significant positivity rates of up to 80% *(n=354)*, followed by 12% *(n=54)* in secondary care hospitals and 8% *(n=33)* from community cohorts. The isolation rates of *S*. Paratyphi A were compared with *S*. Typhi to obtain the proportion that was found to range between 1:5 to 1:11 across various sites, as described in **Suppl Table 3**. Overall, *S*. Paratyphi A comprised 17.6% (441/2503) of total enteric fever cases in India and was majorly recorded in the tertiary care settings.

### Antimicrobial susceptibility testing of S. Paratyphi A isolates

The antimicrobial susceptibility test demonstrated that 100% of *S*. Paratyphi clinical isolates (*n=441*) were non-MDR and susceptible to each of the first-line antibiotics (ampicillin, chloramphenicol, and trimethoprim-sulfamethoxazole). Fluoroquinolone non-susceptibility remained at nearly 98.9%, while a high degree of susceptibility to current alternative treatment options was recorded (100% susceptibility to azithromycin and ceftriaxone) (**Suppl Table 4)**. Overall, Indian *S*. Paratyphi A isolates were found to be generally non-susceptible to ciprofloxacin, while they continue to be susceptible to first-line agents.

### Phylogeny and Population structure of S. Paratyphi A

Phylogenetic relationship of 552 *S*. Paratyphi A isolates based on 4,458 core genome SNPs showed the distribution of study isolates within a global genomic framework. The observed global phylogeny clustered the isolates into seven previously defined lineages (A-G), in which the study isolates were distributed between lineage A (65.8%; 100/152), C (26.3%; 40/152) and F (7.9%; 12/152) (**Figure 1**). RhierBAPS (level 1) yielded five clusters, while level-2 clustering has distinguished a total of 21 sub-lineages (**Suppl Table 2**). Though previous studies have described the sub-lineage level distribution of *S*. Paratyphi A isolates, we have used the recently developed ‘Paratype scheme’ [15] to define sublineages/genotypes within lineage A, C and F. We identified nine genotypes (2.4.1 - 2.4.9) within the dominant lineage A (genotype 2.4) based on Paratype scheme. Geographical distribution of lineage A isolates showed genotype 2.4.3 (previously A1) being predominant in Nepal, 2.4.1 (formerly A2) was present in both Nepal and India and 2.4.4 (previously A3) was primarily found in Bangladesh. Genotype 2.4.2 was predominantly seen in India with a sparse presence in other South Asian countries. The Paratype scheme assigned five new genotypes (2.4.5 – 2.4.9), mainly consisting of Indian isolates. Among the new genotypes, 2.4.5 have been circulating globally, while 2.4.6, 2.4.7 and 2.4.8 consist of Indian isolates distributed distinctly in different geographic regions across the country. Notably, genotype 2.4.9 was geographically confined to a single site in Northern India, indicating a large localized outbreak. The geographical distribution of Paratyphi A genotypes from the study collection is shown as a scattered pie chart (**Suppl Figure 1**).

**Figure 1:**
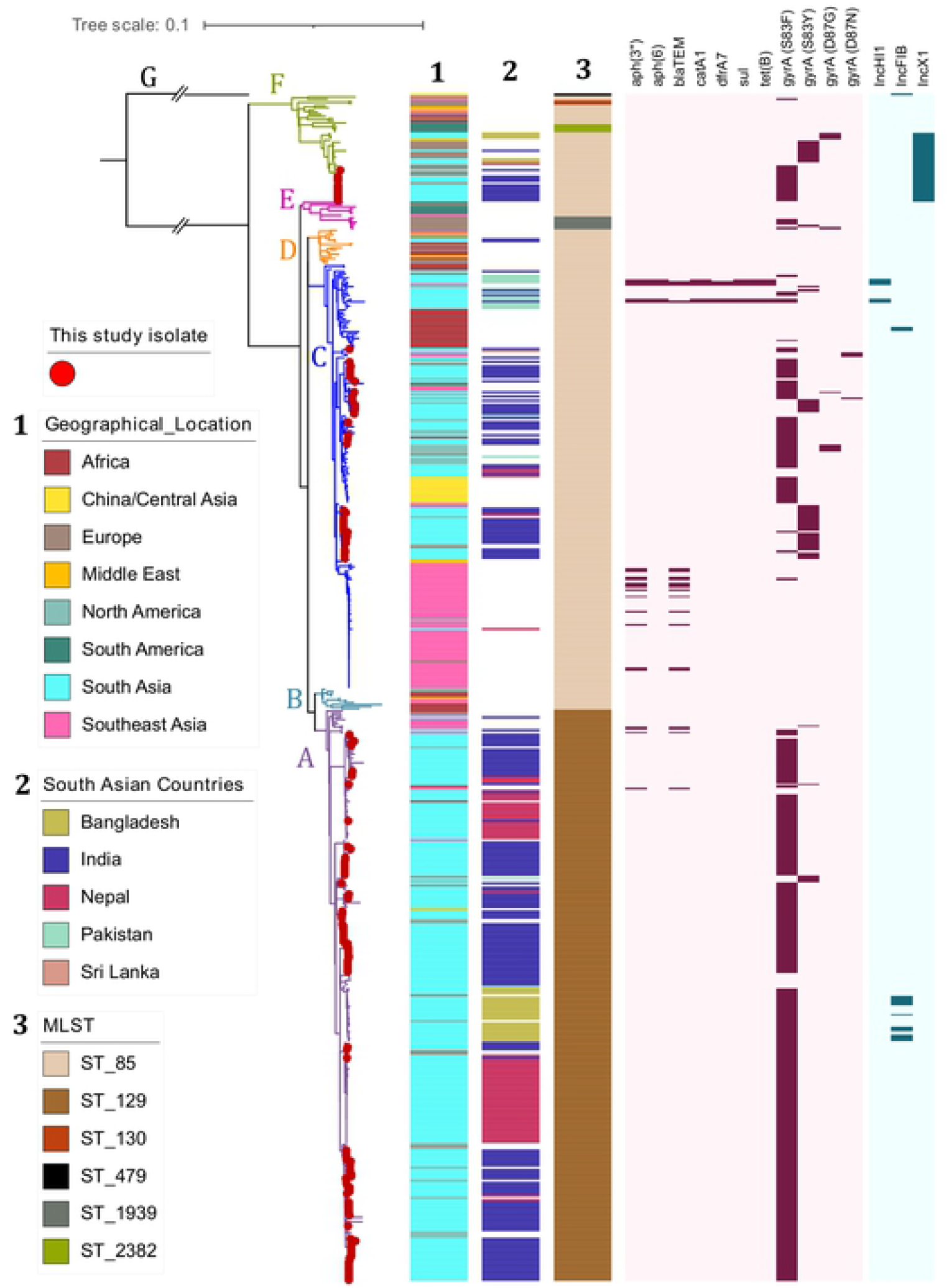
Phylogenetic distribution of contemporary Indian S. Paratyphi A isolates in a global context: Rooted maximum likelihood phylogenetic tree of contemporary Indian *S*. Paratyphi A (*n=152)*, combined with global genome collection (*n=400*) representing the current global distribution. The tree was derived from 4286 SNPs mapped against the reference genome of *S*. Paratyphi ATCC 9150 (Accession No: CP000026.1) using Snippy and rooted to the outgroup strain (ERR028986: Lineage G). Red-colored dots at the tip of the branches indicates the position of this study isolates. Contemporary Indian *S*. Paratyphi A isolates of this study were found distributed across the global tree with both lineages A, C and F. Genomes with their respective metadata are labeled as color strips and key for each variable were mentioned. Strip 1 and 2 indicates the location and 3 represent MLST of each isolate. Heatmap represents the QRDR mutations that confer resistance to fluoroquinolone and presence of plasmids. Scale bar indicates substitutions per site. Color keys for all the variables are given in the inset legend. The tree was visualized and labeled using iTOL (https://itol.embl.de/).

The existing population structure defining sub-lineages of C (C1-C5) was not consistent with rhierBAPS clusters due to its genomic diversity and broad geographical representation, unlike regionally restricted lineage A. Sub-lineages C1 and C2 were represented by polytomies while C4 and C5 were not following the BAPS level 2 clustering (**Suppl Table 2**). The classification of lineage C (2.3) based on the Paratype scheme provides genotypes 2.3.1 (previously C5), 2.3.2 and 2.3.3 (formerly C4). Geographical distribution of global *S*. Paratyphi A isolates showed genotype 2.3 (previously C3) was represented by isolates originating from Africa and Pakistan. Genotype 2.3.2 were isolates predominantly from south Asia, while genotype 2.3.3 isolates were mainly from China, Southeast Asia, and South Asia. Similarly, the first cluster in sub-lineage C5 was designated as genotype 2.3.4 with isolates almost exclusively from India (80%; 20/25), whereas the second cluster (referred to as 2.3.1) was represented by outbreak isolates from Cambodia (**Suppl Figure 2)**. Genotyping of lineage F (genotype 1) was predicted to contain four sub-clusters (1, 1.1, 1.2.1 and 1.2.2), of which 1.2.2 comprised contemporary *S*. Paratyphi A isolates from both India (the present study isolates) and Bangladesh.

### MLST, quinolone resistance mutations and plasmids

Isolates belonging to lineage A were grouped into sequence type 129 (ST129), while lineages B-F were predominantly ST85. The single isolate clustered in lineage G was distinct and belonged to ST479, a double locus variant of ST85. The isolates in our study were pan-susceptible to antibiotics, except for fluoroquinolones. Resistance to the first-line antibiotics (ampicillin, chloramphenicol and co-trimoxazole) was not observed among our study isolates. In contrast, a few *(n=4)* isolates from the global collection were multidrug-resistant (MDR). Genes associated with the MDR phenotype (*bla*_TEM_, *cat, dfrA, sul*) were absent in all study isolates.

Fluoroquinolone non-susceptibility in dominant lineages (A, C and F) of *S*. Paratyphi A was driven mainly by *gyrA-*S83F substitutions, with a few isolates harboring *gyrA-*S83Y (predominantly genotype 2.3) variant. Also, a significant number of isolates were fluoroquinolone susceptible with no mutations in the quinolone-resistance-determining region (QRDR), particularly genotype 2.3.1 (**Figure 1)**. Plasmid profiling revealed that most of the lineage C isolates *(n=116)* harbored a ColRNAI plasmid with no AMR genes. Interestingly, isolates belonging to genotype 1.2.2 *(n=27)* possessed IncX1 plasmid, while the MDR isolates from the global collection carried the AMR genes in either IncFIB or IncH1B plasmid.

### Lineage-specific evolution of S. Paratyphi A

Mutations and gene flux that defines or drives the lineages or sub-lineages of *S*. Paratyphi A were identified from the population structure. The role of gene flux in evolution was determined by pan-genome analysis, while gene inactivation (frameshift mutations) and non-synonymous substitutions were determined by accessing the variant type. Synonymous mutations were not considered as their effect on evolution is likely negligible on the short evolutionary timescale captured in modern molecular epidemiological studies.

Pan-genome analysis revealed the variation of gene content between *S*. Paratyphi A genomes. About 73.8% (3944/5344) were considered core genes (found in >99% genomes), while 18.7% (997) genes were shared by ≤15% of isolates among the 552 screened (**Suppl Fig. 3**). Lineage-specific gene gain or loss during the evolutionary process showed the phylogenetically distinct lineage G lack SPI2. (**Table 1**). Gene gains that likely represent the host adaptation or pathogenicity with respect to the phylogenetic lineages were rather limited to mobile genetic elements. For example, the C4 sub-lineage (genotype 2.3.2 and 2.3.3) of *S*. Paratyphi A has acquired prophage regions P2/ PSP3 phage that could account for their host specificities (**Suppl Fig. 4)**. Interestingly, genotype 1.2.2 was found to have acquired IncX1 plasmid while the plasmid was absent in the older isolates from the lineage F (**Figure 1)**.

**Table 1:** Loss and Gain detected between phylogenetic lineages/genotypes of *S*. Paratyphi A

| S. No | Gene/ Region | Lineage/Genotype | Remarks |
| --- | --- | --- | --- |
| 1 | SPI-2 | A-F | Either lost in G or gained by A-F |
| 2 | P2/ PsP3- like phage | 2.3.2/2.3.3 | Gained by 2.3.2/2.3.3 (C<sub>4</sub>) |
| 3 | IncX1 plasmid | 1.2.2 | Gained by 1.2.2 |

Accumulation of pseudogenes/genome degradation events during the evolution provides insights into the continuous host adaptation or adaptive selection of *S*. Paratyphi A. We identified several lineage-specific pseudogenes since they diverged from ancestral lineages. In addition to the 133 pseudogenes conserved across all lineages except in lineage G, 50 additional genes were identified to be associated with loss of gene function through nonsense substitutions or frameshift mutations (**Suppl Table 6**). A total of 10 pseudogene-forming mutations that could be associated with the emergence of lineages are listed in **Table 2**. Gene flux information and pseudogenes specific to lineages during the evolution of *S*. Paratyphi A are overlaid on a timed phylogenetic tree generated using Figtree in **Figure 2**.

**Table 2:**
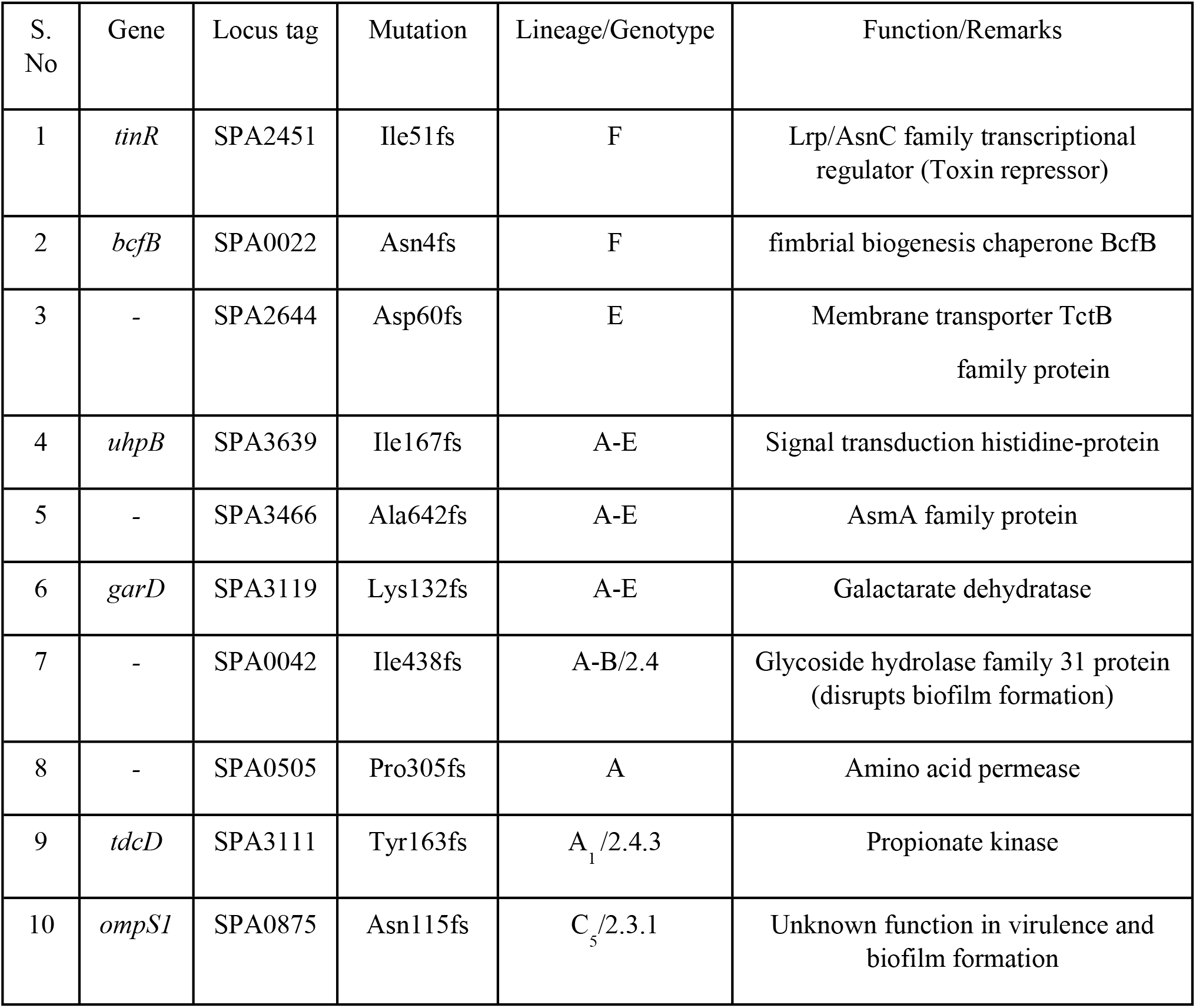
List of functional gene inactivation mutations identified between phylogenetic lineages

**Figure 2:**
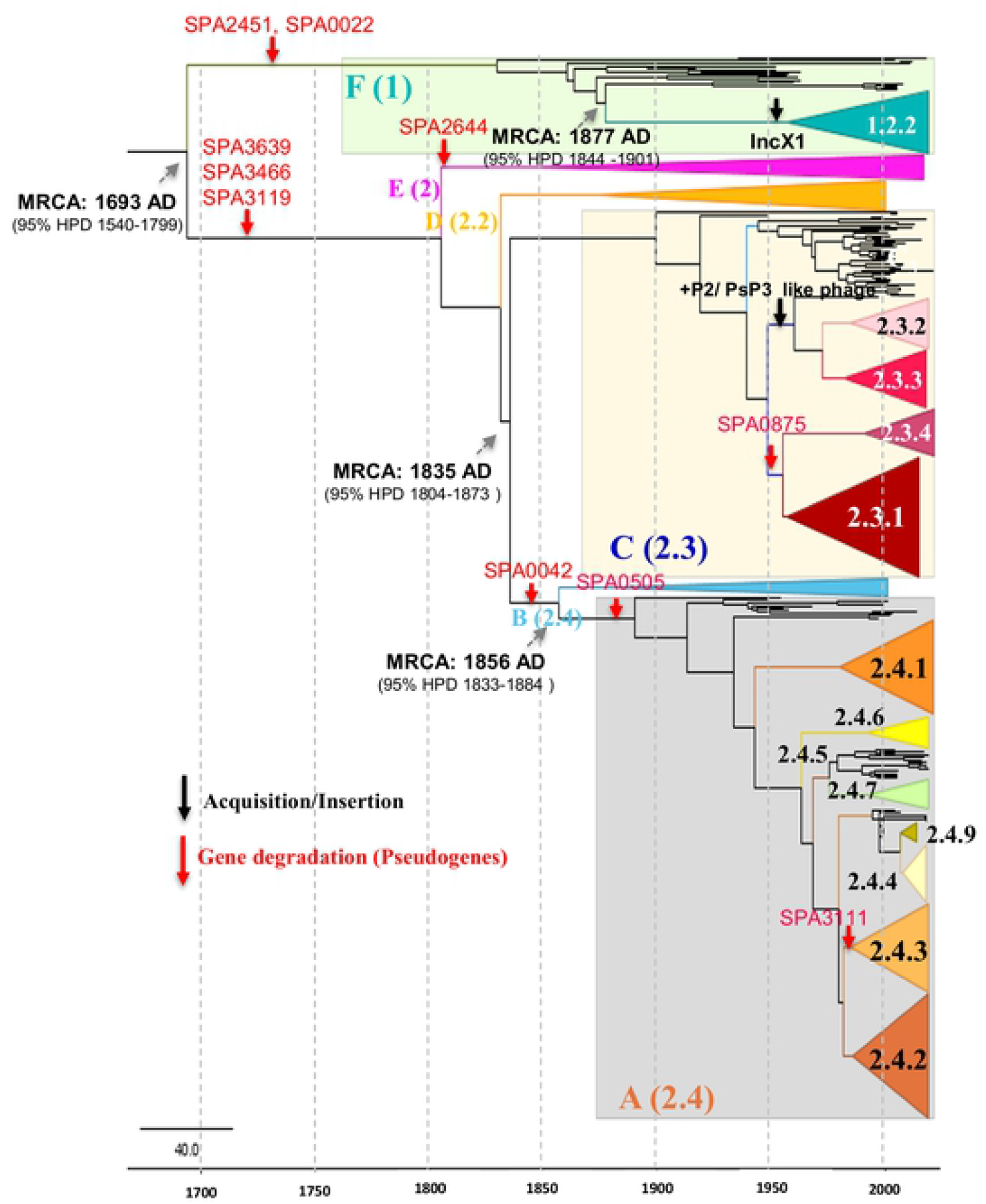
Time-calibrated Bayesian phylogeny phylogenetic tree showing the evolutionary events (pseudogene forming mutations, insertions and deletions) that define the seven modern lineages and sub-lineages of *S*. Paratyphi A. Major lineages/ genotypes were simplified as colored cartoon triangles using FigTree (http://tree.bio.ed.ac.uk/software/figtree/). Red arrow represents frameshift mutation/ gene degradation, Black arrow represent acquisition/ gene gain. Grey arrows demarcate nodes of interest, and the accompanying data indicate 95% HPD of node heights.

Time-scaled Bayesian phylogenetic analysis showed that the model combination best fitted the data was a relaxed molecular clock paired with a constant population size. This analysis dated the most recent common ancestor (MRCA) of *S*. Paratyphi A to the year 1693 (95% HPD 1540-1799) when a single isolate belongs to the distinct clade (lineage G) was excluded for bayesian analysis. The dominant lineages C and A have likely diversified between 1835 (95% HPD: 1804-1873) and 1856 (95% HPD: 1833-1884), respectively. Similarly, genotype 1.2.2 is estimated to have expanded in the year 1877 (95% HPD: 1844 -1901) by acquiring IncX1 plasmid. Overall, the estimated evolutionary rate was was 4.008 × 10^−5^ or 0.301 substitutions/site/year (s/s/y).

### Mutations in O:2-antigen biosynthesis genes

Mutation analysis of O:2-antigen biosynthesis genes (*rfb* region) showed the region carrying six characteristic missense mutations in comparison with ATCC9150 reference strain (vaccine candidate). Mutations in *rfb* gene cluster consist of single amino acid substitutions in *rfbG* (H348R), *rfbD* (G262S), *rfbE* (S167L), *rfbS* (C249S), *rfbB* (H176Y) and *rfbC* (E154K). Interestingly, these mutations are possibly associated with positive selection of lineages/genotypes currently circulating in south Asian countries (**Suppl Fig. 5**). For instance, genotypes carrying a characteristic missense mutation in LPS O-antigen biosyntheses such as 1.2.2 (*rfbC*: E154K), 2.3.3 (*rfbS*: C249S) and 2.4.2 (*rfbD*: G262S) are increasingly being detected, particularly in India. The impact of these lineage-specific mutations on the protein structure is still unknown, although slide agglutination tests showed no significant difference between mutant genotypes and the wild-type strain. The predicted free energy gap difference (ΔΔG) between the wild type and mutant protein measures how the mutation impacts the protein stability. The ΔΔG values of different *rfb* gene mutations indicated stabilizing scores except for *rfbC*: E154K (**Suppl Table 7)**. However, the significance of these mutations in the LPS structure and the potential impact on current vaccine development is yet to be studied.

## Discussion

Genome analysis of 152 *S*. Paratyphi A isolates collected from different geographical locations in India between 2017-2020 revealed evolutionary changes that favor genetic diversity for its persistence and spread. Comparative genome analysis unambiguously placed the contemporary *S*. Paratyphi A isolates from India into three lineages, with lineages A and C being dominant. This concurs with the previous analysis that reported the placement of present-day south Asian isolates in these three lineages [9,12,16]. Further extension of the current designation of sub-lineages that belong to lineages A, C and F based on the recently developed Paratype genotyping scheme [15] has improved the sub-lineage level classification of major lineages. Our results provide a more detailed picture of the population structure and geographical distribution of *S*. Paratyphi A isolates in south Asian countries, particularly in India. Overall the contemporary Indian *S*. Paratyphi A isolates clustered closely with isolates originating from Bangladesh, Nepal and Pakistan, suggesting the regional circulation of these lineages across south Asia.

Geographical distribution of genotypes confirms the dominance of *S*. Paratyphi A isolates from Nepal (2.4.3 & 2.4.1), Bangladesh (2.4..4) and India (2.4 and 2.4.2) in the sub-clusters of A. Within lineage C, genotype 2.3 predominantly contains isolates from Africa and Pakistan. Similarly, isolates from India (2.3.2 & 2.3), China (2.3.3) and Cambodia (2.3.1) were distributed as geographically confined sub-lineages, respectively [17,18]. The phylogenetic positioning of contemporary *S*. Paratyphi A isolates in lineage F was unexpected; however, recent reports from Bangladesh also documented similar findings [15,16]. A closer look at the lineage F isolates revealed the positioning of older isolates from the global collection in genotype 1/1.1, while the contemporary isolates from India and Bangladesh form the genotype 1.2.2. The emergence of genotype 1.2.2 can be attributed to the acquisition of IncX1 plasmid, highlighting the role of horizontal gene transfer in favoring the successful evolution and long-term persistence of these clones.

Antimicrobial resistance determined by phenotypic and genomic analysis of the study isolates showed low-level resistance to antimicrobials except for fluoroquinolones. These results were consistent with the previous estimates as most of the studies from south Asia report either no or low levels of multidrug resistance [19]. Though MDR phenotypes were observed in a few *S*. Paratyphi A strains from the global collection, the plasmid was eventually lost during the evolution due to the greater fitness of antibiotic-sensitive strains [12]. On the contrary, fluoroquinolone non-susceptibility (FQNS) was high amongst *S*. Paratyphi A in South Asia, with FQNS strains from the SEFI collection accounting for 98% of all isolates [20].

The FQNS *S*. Paratyphi A were predominantly single QRDR mutant (*gyrA*-S83F) and distributed across the dominant phylogenetic lineages (A, C and F). Interestingly, the successes of all three lineages/sub-lineages in south Asian countries appear to be largely driven by the development of *gyrA* S83F mutation (except for a subcluster in 2.3 -*gyrA* S83Y). Though this mutation is not unique to these lineages, there is a strong association between reduced susceptibility to fluoroquinolones caused by the S83F mutation and the persistence/spread of these lineages. Our data is in line with the emergence of FQNS *S*. Typhi lineages with positively selected S83F mutant in south Asian countries [21]. Nevertheless, acquired AMR genes or mutations within these QRDR regions are not the sole factors that determine the evolution of *S*. Paratyphi A [18].

The evolution of *Salmonella* sp. is strongly associated with gene influx, genome degradation and rearrangement events that aid in host adaptation [22]. Modern isolates of *S*. Paratyphi A possess an average of 173 genome degradation events through pseudogene formation in comparison to the 25-35 pseudogenes observed in host generalists, such as *S*. Typhimurium [23]. Since *S*. Paratyphi A evolved into a human-specific systemic pathogen approximately 450 years ago, many of these adaptive mutations would have occurred very early [12]. The genetic features responsible for causing enteric fever were a perpetual change, while the recent microevolution is transient and will likely be removed by purifying selection in the future [12].

In our study, we also focused on critical events that may have contributed to the expansion or extinction of the seven modern lineages of *S*. Paratyphi A. Our observations indicate that the emergence of these lineages and sub-lineages was primarily associated with gene acquisitions or losses and mutations in genomic regions related to metabolism (**Fig. 2**). Pan-genome analysis of SEFI isolates and representative isolates from a global collection showed the gain of prophages or plasmids during the selection of lineages (**Table 1)**. Evaluation of gene degradation also depicted that disruption of metabolic pathways along the phylogenetic lineages/sub-lineages are key factors in evolution (**Table 2)**. These findings further confirm that differences in metabolic functions due to environmental and/or human behavioral factors play a significant role in the expansion of lineages.

Identifying missense mutations occurring specifically in genes responsible for LPS biosynthesis is crucial since these genes are the critical targets for developing vaccines and diagnostic assays [24]. Though the impact of these mutations on phenotype, fitness and evolution is currently unknown, the presence of lineage/genotype-specific association may be considered as a signature of positive selection [25]. Among the six missense mutations, at least five have been predicted to stabilize the protein structure (ΔΔG ≥ 0). Serotyping the genotypes (carrying *rfb* loci mutations) by slide agglutination confirmed good agglutination with the O2 antisera, which suggests no or low impact structural changes in LPS. However, the experimental impact of these mutations will require more laboratory analyses. Further sequencing of isolates may reveal the existence of any selective pressure that may aid the genotypes in evading the host immune response. At present, the *S*. Paratyphi A O-polysaccharide glycoconjugate vaccine will have a protective response against all currently circulating *S*. Paratyphi A lineages.

Several isolates belonging to the global collection could not be assigned to genotypes by Paratype, which would require sequencing of more *S*. Paratyphi A isolates from the region in the future. We could robustly evaluate the global phylogenomics of this mostly neglected pathogen with the collection we had. Still, more extensive studies and continuous surveillance is needed to draw better public health policies for *S*. Paratyphi A control.

## Materials and Methods

### Study settings

A total of 19 centers across the country, with a diverse and vast population, in a three-tiered surveillance system consisting of community-level health care setting (Tier 1), secondary hospitals (Tier 2) and tertiary care hospitals (Tier 3) were selected to form an Indian Typhoid network entitled “Surveillance of Enteric Fever in India” (SEFI) [26]. Details of the isolates, participation centers and respective epidemiological settings are provided in the supplementary material (**Suppl Table 1**).

### Bacterial isolates and antimicrobial susceptibility testing

Clinical isolates of *S*. Paratyphi A isolated from blood and bone marrow cultures from the participating centers were received at the central reference laboratory at the Department of Clinical Microbiology, Christian Medical College, Vellore, India. These isolates were further identified and confirmed as *S*. Paratyphi A by standard biochemical and agglutination tests by the Kauffmann*-*White scheme [27]. Antimicrobial susceptibility testing was performed for the commonly used agents such as ampicillin (10 μg), chloramphenicol (30 μg), co-trimoxazole (1.25/23.75 μg), ciprofloxacin (5 μg), pefloxacin (5 μg), ceftriaxone (30 μg) and azithromycin (15 μg) by disk diffusion. Test results were interpreted as per clinical breakpoints recommended by the Clinical and Laboratory Standards Institute [28]. Azithromycin zone size interpretation was based on CLSI *S*. Typhi criteria (Sensitive ≥13 mm; Resistant ≤12 mm)

### Genomic DNA extraction and Sequencing

A subset of 152 *S*. Paratyphi A isolates from the collection (*n=152)* were selected for WGS by ensuring temporal and geographic representation across India. Each bacterial isolate was grown in LB broth (Oxoid) at 37°C and growth was assessed by the increase in turbidity and by microbial count (>10^9^ cfu/ml). The liquid cultures were centrifuged at 10,000 rpm and DNA was extracted from the pelleted cells using Wizard DNA purification kit (Promega, Madison, USA) as per the manufacturer’s protocol. The purity and concentration of extracted DNA were measured using Nanodrop One (Thermo scientific) and Qubit dsDNA HS Assay Kit (Life Technologies).

Sequencing ready, paired-end library was prepared using 100 ng of DNA with the Nextera DNA sample preparation kit as per the manufacturer’s instructions (Illumina, Inc., San Diego, USA). This was followed by sequencing on Illumina NextSeq 500 and HiSeq X 10 platforms with a paired-end run of 2×150 bp. Raw reads were quality checked to remove adapters and the filtered high-quality reads were assembled using Unicycler (https://github.com/rrwick/Unicycler).

### Genome data acquisition and characterization

A global representation of *S*. Paratyphi A *(n=400)* isolates was selected from a curated subset of Enterobase (http://enterobase.warwick.ac.uk/species/senterica/) and other previously published genomes [9, 12, 13, 16 – 18]. The corresponding paired-end reads were downloaded from European Nucleotide Archive (ENA; http://www.ebi.ac.uk/ena). <u>Genotypes were assigned from raw reads using Paratype</u> (https://github.com/CHRF-Genomics/Paratype). The high coverage (>50X) reads were assembled using Unicycler v0.4.9 (https://github.com/rrwick/Unicycler). The assembled genomes were analyzed using Seqsero v2.0 [29] to confirm the antigenic profile of the serotype. Sequence types of the isolates were designated using the Multilocus sequence typing (MLST) pipeline available in the Center for Genomic Epidemiology (CGE) (https://cge.cbs.dtu.dk/services/). AMR genes, point mutations and plasmids were screened against resfinder and PlasmidFinder database by using ABRicate (https://github.com/tseemann/abricate). In total, 152 *S*. Paratyphi A study isolates from SEFI collection along with 400 genome sequences from the public database were included. The complete list of genomes used in this study and metadata is available in **Suppl Table 2**.

### Variant calling and Phylogenetic Tree construction

The assembled genomes were mapped against the reference genome *S*. Paratyphi A ATCC 9150 (Accession No: CP000026.1) using Snippy v4.6.0 [30]. The core genome SNP differences between the genomes, with respect to the reference, were generated as an alignment file. Further, Gubbins (v.2.3.1) was used to remove the recombination regions from the core genome alignment to produce a recombination filtered alignment file [31]. The Maximum likelihood (ML) phylogenies were constructed using the Fasttree [32] with GTRGAMMA model and the generated phylogenetic tree was visualized and annotated using iTOL [33]. Phylogenetic clusters were assigned using rhierBAPS [34] specifying two cluster levels with 30 initial clusters (snp.matrix, max.depth = 2, n.pops = 30, n.extra.rounds = Inf, quiet = TRUE).

To assess the temporal structure, root-to-tip genetic distances from (ML) tree against sample collection dates using TempEst v 1.5.1 (http://tree.bio.ed.ac.uk) was performed. Using the regression analysis of root-to-tip distances, an association between sampling times and genetic divergence (molecular clock) was determined. The timed evolution of *S*. Paratyphi A lineages was estimated using Bayesian phylogenetic methods available in BEAST v.1.10 [35, 36]. The recombination free alignment file was used as the input for the time-scaled phylogenetic analysis. The Hasegawa, Kishino and Yano model (HKY) substitution with different demographic models (Bayesian skyline, exponential and constant) was investigated. To determine the best-fitting coalescent model to describe changes in effective population size over time, log marginal likelihoods were calculated using path sampling and stepping stone sampling methods. Finally, Bayes factor [37] was used to determine the best fit model with the formula [logBF = logPr(D|M1) – logPr(D|M2)]. The selected bayesian skyline with uncorrelated relaxed clock model was run in 3 independent chains for 200 million with a sampling of 10000 generations. A burn-in of 20% was discarded from each run and resulting log files were combined using LogCombiner 1.8.1 [38]. The convergence and mixing were manually inspected using Tracer.v.1.7 [39] to ensure that all the parameters converged to an ESS of >200. The maximum clade credibility (MCC) tree was generated using Treeannotator v.1.8.2 [40]. The output was analyzed using Tracer v1.7, with uncertainty in parameter estimates reflected as the 95% highest probability density (HPD). The annotated phylogenetic tree was visualized using FigTree v.1.4.4 [41].

### Lineage wise mutation profiling

Mutations were identified by *in-silico* determination of single nucleotide polymorphisms (SNPs) using the Snippy v4.6.0 mapping and variant calling pipeline (https://github.com/tseemann/snippy). To obtain SNPs, the draft genome of the study population was mapped against the annotated feature of reference genome *S*. Paratyphi A ATCC 9150 (CP000026.1). In-house written bash scripts were used to retrieve the pattern of mutation accumulation with respect to the phylogenetic lineages. Genes that contained either frameshift mutation or a premature stop codon were manually curated and classified hypothetically disrupted coding sequences (HDCS) or pseudogenes. The identified pseudogenes in different lineages were compared with the data reported previously [13,23].

### Pan-genome analysis

The pan-genome of all the study isolates of *S*. Paratyphi A (*n=552*) was annotated using Prokka v. 1.14 [42] using a custom database created with “prokka-genbank_to_fasta_db” based on 1328 annotated *S*. Paratyphi A genomes downloaded from NCBI (https://www.ncbi.nlm.nih.gov/genome/browse/#!/prokaryotes/152/). To remove redundancy, CD-HIT version 4.8.1 was used with the following parameters: -T 0 -M 0 -g 1 -s 0.8 -c 0.90 [43]. The Prokka-compatible protein sequence fasta file (custom database) was confirmed to be used by the Prokka with relevant flags as follows --genus spa --usegenus --rfam --evalue 1e-05 --coverage 50 (https://github.com/tseemann/prokka). The annotated draft assemblies in GFF3 format was used as input to evaluate pan-genome diversity using Panaroo [44]. Panaroo was run using its “strict” mode with ‘remove invalid genes enabled -I option *.gff -o results --clean-mode strict --remove-invalid-genes --core_threshold 0.98 -t 6 -c 0.80. The gene presence or absence in each genome obtained were grouped according to the phylogenetic lineages (A-G) using twilight scripts (https://github.com/ghoresh11/twilight) with default parameters [45]. Gene gain or loss was curated manually and mapped into the timed Bayesian phylogenetic tree generated using Figtree (http://tree.bio.ed.ac.uk/software/figtree/).

### Mutations in LPS biosynthesis genes

Snippy based variant calling was performed on the assembled genomes (*n=551*) using the *rfb* loci of strain ATCC9150 (CP000026: 860063 – 884690) as the reference. SNPs and Indels occurring within the coding region of *rfb* loci were considered and the mutations were screened and arranged according to phylogenetic lineage in tabulated format. Whole-genome alignment (.full.aln) from the snippy output was used to build a maximum likelihood phylogeny using FastTree [32] with GTRGAMMA model. The generated phylogenetic tree was visualized and annotated using iTOL. The three-dimensional structures of rfb genes were modelled using ModWeb (https://modbase.compbio.ucsf.edu/modweb/) homology-based method. The quality of the model was evaluated using Ramachandran plot and the effect of mutations at a molecular level were then further analyzed using FoldX version 4 (http://foldxsuite.crg.eu/node/196).

## Data availability

Whole genome sequenced raw read data is available at the European Nucleotide Archive (ENA) and individual sample accession numbers are listed in Supplementary Table S2

## Acknowledgments

We thank Prof. Nicholas Grassly, Imperial College London, for assistance with study design and research proposal development and, Arif M. Tanmoy, CHRF, Dhaka for help with the genotyping analysis. We acknowledge Dr. Duncan Steele, Ms. Megan Carey & Dr. Supriya Kumar, Bill & Melinda Gates Foundation for their technical support throughout the study on behalf of SEFI consortium. We thank all the lab members involved in SEFI reference lab activities, especially Dr. Anushree Amladi, Ms. Baby Abirami S, Ms. Dhanabhagyam K, Ms. Beebi E, Ms. Suganya S, Ms. Udaya and Mr. Ayyanraj N, CMC Vellore implicated in phenotypic testing and stock culture maintenance. We would also like to thank all the members of SEFI consortium, Wellcome Trust Research Laboratory, CMC Vellore and core sequencing teams at the Wellcome Trust Sanger Institute for their contribution to genome sequencing. The authors thank Ms. Catherine Trueman (Clinical Pharmacist, CMC Vellore) for helping with language editing. This study was approved by the Institutional Review Board (IRB) of Christian Medical College, Vellore (IRB Min No: 10393 dated 30.11.2016).

## Supporting Information

**Suppl Fig. 1**: Map of India showing the regional diversity of *S*. Paratyphi A genotypes. Pie chart colours indicate the propotion of genotypes prevalent in three major geographical locations in India. Study sites are represented as per the settings. Color keys for all the variables are given in the inset legend

**Suppl Fig. 2:** Rooted maximum likelihood phylogenetic tree of *S*. Paratyphi A isolates showing the comparative phylogenetic clustering by lineages, predefined sub-lineages, RhierBAPS population clustering (level 1) and Paratype genotyping scheme. Lineages are represented by various colored branches. Sublineages, BAPS cluster and Paratype scheme are labeled as color strips.

**Suppl Fig. 3:** Visualization of pan-genome analysis data by Panaroo of 552 *S*. Paratyphi A genomes. (a) Pie chart indicates the core, soft core, shell and cloud genome composition of *S*. Paratyphi A genomes (b) Maximum likelihood tree of *S*. Paratyphi A genomes were compared to a matrix with the presence (blue) and absence (white) of the accessory genes found in the pan-genome. The image was prepared using Phandango (https://jameshadfield.github.io/phandango/#/)

**Suppl Fig. 4:** Linear representation of acquired prophage regions (P2/ PSP3 phage) generated using Proksee (https://proksee.ca/) available at the CG view server (https://cgview.ca/)

**Suppl Fig. 5:** Rooted maximum likelihood phylogenetic tree of *rfb* loci of *S*. Paratyphi A isolates derived from the whole genome alignment by mapping against the reference genome of *S*. Paratyphi ATCC 9150 (Accession No: CP000026.1) using Snippy. Lineages and genotypes are labeled as color strips. Amino acid substitutions in the *rfb* loci are represented by heatmaps.

**Suppl Table 1:** List of whole genome sequenced isolates collected from the participating sites of SEFI network

**Suppl Table 2:** List of *S*. Paratyphi A genomes used in this study with accession IDs and metadata

**Suppl Table 3:** Distribution of *S*. Typhi and *S*. Paratyphi A isolates collected across the participating sites of SEFI network

**Suppl Table 4:** Antimicrobial susceptibility profile of *S*. Paratyphi A tested in the present study

**Suppl Table 5:** Lineage-defining Frameshift mutations/stop codons in *S*. Paratyphi A genomes

**Suppl Table 6:** Lineage-defining missense mutations in *S*. Paratyphi A genomes

**Suppl Table 7:** List of lineage defining mutations in the O:2-antigen biosynthesis genes (*rfb* region) of *S*. Paratyphi A and their predicted impact on protein structures

